# Towards Unmanned Proteomics Data Generation: A Fully Automated Sample-to-Data System for Proteomic Experiments

**DOI:** 10.1101/2025.06.02.656978

**Authors:** Dongxue Wang, Wendong Chen, Linhai Xie, Ying Xu, Chuanxi Huang, Yuanyuan Liu, Xi Wang, Xiaowei Huang, Keren Zhang, Mengting Pan, Shaozhen Wang, Baoyi Qin, Yun Yang, Jing Yang, Liujun Tang, Ruijun Tian, Fuchu He

## Abstract

The π-HuB project requires proteomic measurements of millions of low-input samples that are challenging with current technology. Here, we introduce a pilot π-HuB data factory that features a fully automated sample-to-data system to enable high-throughput digitization of human samples. Our platform, as a proof-of-concept towards unmanned proteomics, was demonstrated to show longitudinal robustness and applied for spatial proteomics and the profiling of plasma and cell lines, thereby serving as the workhorse to drive π-HuB.

## Main text

Data-driven biology requires comprehensive and high-quality data at all molecular levels. LC-MS-based proteomics has emerged as the leading method for comprehensively identifying and quantifying proteins in biological samples^1^. Many studies have highlighted the value of having large-scale MS-based proteomic datasets (i.e., from hundreds to more than one thousand measurements) to promote our understanding of human biology and various complex diseases. These efforts have therefore inspired the international community to launch the π-HuB (The Proteomic Navigator of the Human Body) project^2^, aiming to transform biology and medicine in the way that the Human Genome Project did^3^.

Same as with other data-driven big-science projects, π-HuB will start by manufacturing enormous data. Although up to 10,000 proteins could be profiled in a single LC-MS/MS run^4, 5^, paralleling the depth made by state-of-the-art genome sequencing, a central challenge is to raise throughput and data robustness in proteomic measurements without compromising the depth of identification and the quantitative precision^6, 7^. In the past few years, we have witnessed many technical advances in high-throughput proteomics in terms of automated sample preparation and LC-MS/MS technologies. For example, automated sample processing with liquid-handling devices allows the preparation of hundreds of low-input samples per day.^8–10^ In addition, fast chromatographic separations and MS/MS data acquisitions have been achieved by implementing new instruments and/or settings, allowing the analytical throughput of >100 samples per day (SPD) on a single LC-MS instrumentation^11^ ^12^.

Despite significant advancements, the demands of the π-HuB for generating data from millions of samples highlight an urgent need to improve the efficiency and reliability of proteomics data production. First, there is a necessity for a fully automated sample-to-data system that operates without human intervention, eliminating errors or variabilities caused by personnel or batch differences. Second, achieving ultra-high-throughput data generation is essential to reduce the turnaround time from sample preparation to data processing. Lastly, increasing throughput in proteomic measurements should not compromise the depth of profiling or quantitative precision. To address all these needs simultaneously, we present the pilot π-HuB data factory, which features a fully automated sample-to-data system, referred to as the π-Station, designed to facilitate high-throughput proteomics in a nearly unmanned manner **(Supplementary Video 1)**.

Previous automation efforts in LC-MS/MS-based proteomics have primarily concentrated on sample preparation with manual assistance ^13^. The entire process, from sample to data, still requires significant hands-on experience from skilled scientists (**Fig.1a, Extended Data Fig. 1a and b**). Therefore, we designed and set up the π-Station that can seamlessly integrate fully automated sample preparation with LC-MS/MS instrumentation and computing servers, enabling the direct generation of protein quantification data matrices from biospecimen samples without manual intervention (**Fig. 1b**). The π-Station is a linear, expandable, and connectable platform, consisting of 14 customized devices from 7 different vendors, and is seamlessly connected to LC-MS/MS systems via 2 robotic arms. All these hardware are configured in and managed by the Momentum Workflow Scheduling Software, enabling modular method editing of both sample preparation and LC-MS/MS data acquisition (**Extended Data Fig.2a**). Meanwhile, data storage, processing, quality control, and monitoring modules can be achieved by controlling external proteomic software tools and social media applications via in-house UI tools developed in Python.

**Fig.1.**
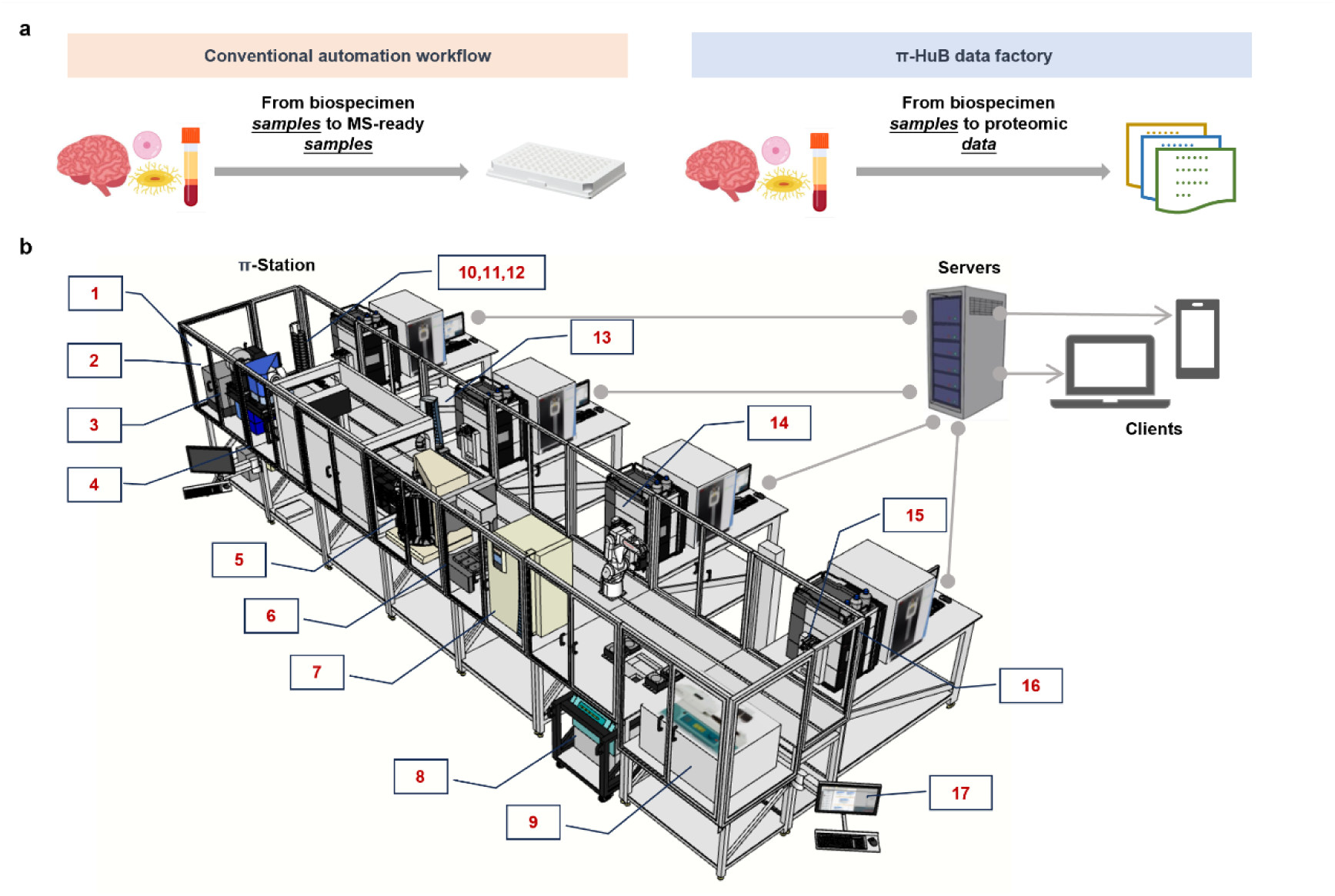
Overview of the π-HuB data factory. **a,** Comparison of the conventional automated proteomic workflow and our sample-to-data fully automated workflow. **b,** Overview of the prototype of π-HuB data factory. It mainly consists of π-Station, LC-MS/MS instrumentation, and computing servers. π-Station is the fully automated sample preparation module, which is coupled to the LC-MS/MS systems. This prototype includes the following hardware and software: (1) LE220Rsc focused-ultrasonicator, Covaris; (2) ALPS 3000 automated microplate heat sealer,Thermo Fisher Scientific; (3) XPeel automated plate seal remover, Brooks;(4) Biomek i7 automated liquid handler with the Amplius positive pressure extractor, Beckman Coulter; (5) Cytomat 10 hotel, Thermo Fisher Scientific; (6) AssayMAP Bravo Protein Sample Prep Platform, Agilent; (7) Cytomat 2C425 automated incubator, Thermo Fisher Scientific; (8) Rotanta 460 automated centrifuge, Hettich; (9) CombiDancer vortex vacuum concentrator, Hettich; (10) Multiskan SkyHigh microplate spectrophotometer, Thermo Fisher Scientific; (11) Automated Thermal Cycler (ATC), Thermo Fisher Scientific; (12) Incubator Shaker DWP, Inheco; (13) Spinnaker microplate robot, Thermo Fisher Scientific; (14) F7 Industrial Robot, Thermo Fisher Scientific; (15) Vanquish UHPLC Loader, Thermo Fisher Scientific; (16) LC-MS system, Vanquish Neo UHPLC system coupled to Orbitrap Exploris 480 mass spectrometer, Thermo Fisher Scientific; (17) Momentum workflow scheduling software, Thermo Fisher Scientific.

At the π-Station, a typical proteomic workflow includes protein extraction, digestion, desalting, solvent evaporation, resuspension of dried MS-ready peptide, and initiation of LC-MS/MS analysis. The devices utilized to perform these steps are described in the extended data section (**Extended Data Fig. 2b)**. With the Momentum software, a workflow can be executed both individually and efficiently in multiple iterations. For example, a single iteration takes 7.5 h to process 96 samples into MS-ready peptides, while it only takes 1.5 h for the second and the following iterations, resulting in finishing 384 samples in 4 iterations within 12 h (**Extended Data Fig.6**). Supplementing labware and reagents allows for infinite iterations of the whole process. Once the MS-ready peptides are made, the Momentum software will instruct robotic arms to send them to the sample loaders to initiate LC-MS/MS data acquisition on multiple instruments. In the pilot π-HuB data factory, for instance, the analytical capacity of the π-Station can be further enhanced by paralleling data acquisition on 15 LC-MS/MS instruments, which greatly increases turnaround rates for digitizing biospecimen samples into proteomic data. For example, the platform enables the highest throughput of 360 SPD using 60-min standard gradients, so those samples automatically prepared from 4-iteration settings can be measured within nearly a single day.

The ultra-high-throughput sample-to-data workflow enabled by the π-Station poses new challenges for continuously monitoring system performance that requires extensive manual inspection by experienced specialists. To address this bottleneck, in addition to implementing an effective quality control (QC) strategy (**Extended Data Fig. 4**), we developed a computational framework called π-ProteomicInfo for automating data storage, processing LC-MS/MS data, and monitoring the performance and status of LC-MS/MS instruments (**Extended Data Fig. 3a**). The process starts with the monitor module, a standalone program installed on computers that control LC-MS/MS systems. This module monitors instrument status (e.g., warnings and errors) and automatically transfers raw data files once each run is completed. After the data transfer, the analyzer module is triggered to generate qualitative and quantitative profiles for each run and all experiment runs, and the QC module initiates extracting QC metrics for data quality assessments (see **Methods** for details). If QC data is found to be unqualified, the controller module will immediately stop data acquisition by sending commands to the instrument. This action helps prevent the irretrievable loss of precious clinical biopsies typically with very limited amounts. Meanwhile, the specialists operating the corresponding instruments will receive notifications via text messages about the QC results and instrument status, allowing them to perform maintenance as soon as possible (**Extended Data Fig. 3b-d**). This framework thus enables automatic feedback control of our platform’s performance and significantly reduces the data processing time.

To evaluate the π-Station, we conducted DIA analyses of HEK 293T cells with protein inputs ranging from 500ng to 20 μg (**Extended Data Fig. 7**). We quantified 3,175 ± 160 (Mean ± SD, the same as below) proteins with the high-throughput LC-MS/MS method and 8,144 ± 429 with the sensitive LC-MS/MS method from 500ng cell lysates. The depth and stability of analysis improved as the protein input increased. We quantified over 7,500 proteins with median coefficients of variation (CVs) less than 8% from 5 μg samples using both high-throughput and sensitive 1 h LC-MS/MS analysis. To evaluate the long-term stability of this platform, we analyzed 10 different cell line samples across 63 days in a total of 18 iterations using HEK 293T cell lysates as quality control (QC). In each iteration, 6 QC samples were randomly placed on the 96-well plate. The variation in protein and precursor identification of QC samples across each iteration, intra-day iterations, and throughout the two-month period remained below 3% and 6%, respectively, with the maximum median CV of protein abundance remaining under 8% (**Extended Data Fig. 8**). The robustness of our platform was further demonstrated by analyzing ten widely used cell lines. On average, we quantified 7,770 to 8,400 proteins across 22 to 720 biological replicates for each cell line, with a maximum median coefficient of variation (CV) of 10.35%. These data depicted the biological features and differences across multiple cell lines (**Extended Data Fig. 9**). Together, these benchmarking results demonstrate the throughput and robustness of this pilot data factory for π-HuB, which is primarily facilitated by the fully automated sample-to-data system and parallel data acquisition.

Next, we sought to demonstrate the utility of our platform for human plasma profiling. The plasma sample preparation workflow was similar to that of cell lines, allowing us to process 96 samples in 7.5 hours and 384 samples in 12 hours and achieve infinite iterations by supplementing labware and reagents. We implemented a 48 SPD DIA method to analyze naïve plasma samples from a clinical cohort of 398 patients diagnosed with liver cancer. The samples were processed over 5 iterations and analyzed within two weeks, with π-ProteomicInfo overseeing these processes. In each iteration, 6 healthy plasma samples were included as the QC. The analysis of the 28 QC samples showed the high reproducibility of our platform: 5,797 ± 73 precursors, 238 ± 12 proteins, median CV of 4.36% inter-iterations. As a result, we quantified an average of 497 proteins in this liver cancer cohort, and there were no obvious batch effects between different iterations (Extended Data Fig. 10). It is important to note that, in this case study, all MS data were collected in standard DIA settings. When coupled with more advanced DIA methods and/or MS instruments ^11, 14^, it could be foreseen that both the depth and throughput of our platform for plasma proteome profiling would further increase.

Encouraged by these results, we integrated SISPTOT (Simple and Integrated Spin-Tip-Based Proteomics Technology), a miniaturized kit for low-input sample preparation^15^, to further evaluate our capability in spatial proteomic analysis (**Fig. 2a**). The workflow established on the π-Station platform allows for the processing of two iterations of sample preparation, totaling 192 laser capture microdissection (LCM) samples every 5.5 hours, and can operate in infinite mode. For quality control, 6 aliquots of 250 ng of mouse brain lysates were used as QC samples, prepared and analyzed alongside tissue slices in each iteration. As a result, the analysis achieved low intra- and inter-iteration CV for protein abundance values (median CV at 13.9%, n = 14) and reproducible quantification of precursors (65,806 ± 3,737) and proteins (5,740 ± 117). Moreover, we generated a spatially resolved mouse brain proteome atlas of a 12μm coronal section at the resolution of 500μm⊆500μm. Proteomic profiling by 60 min DIA measurement resulted in the identification and quantification of a total of 7,300 protein groups in 184 samples with an average of 5,814 (± 378 SD) protein groups per sample at an FDR of < 1% (**Fig. 2b**). Notably, our analysis correctly recapitulated the expected distributions of known protein markers in various anatomical regions (**Fig. 2c**). For example, Cacng3, Slc30a3, and Synpo in the cortex, Ahi1, Baiap3, and Scg2 in the hypothalamus, Ccdc136, Prkcd, and Synpo2 in the thalamus, Ahi1, Baiap3, and Scg2 in the hypothalamus, agreed well with their previously-reported distributions. More examples of regionally enriched proteins are shown in **Extended Data Fig. 5**.

**Fig.2.**
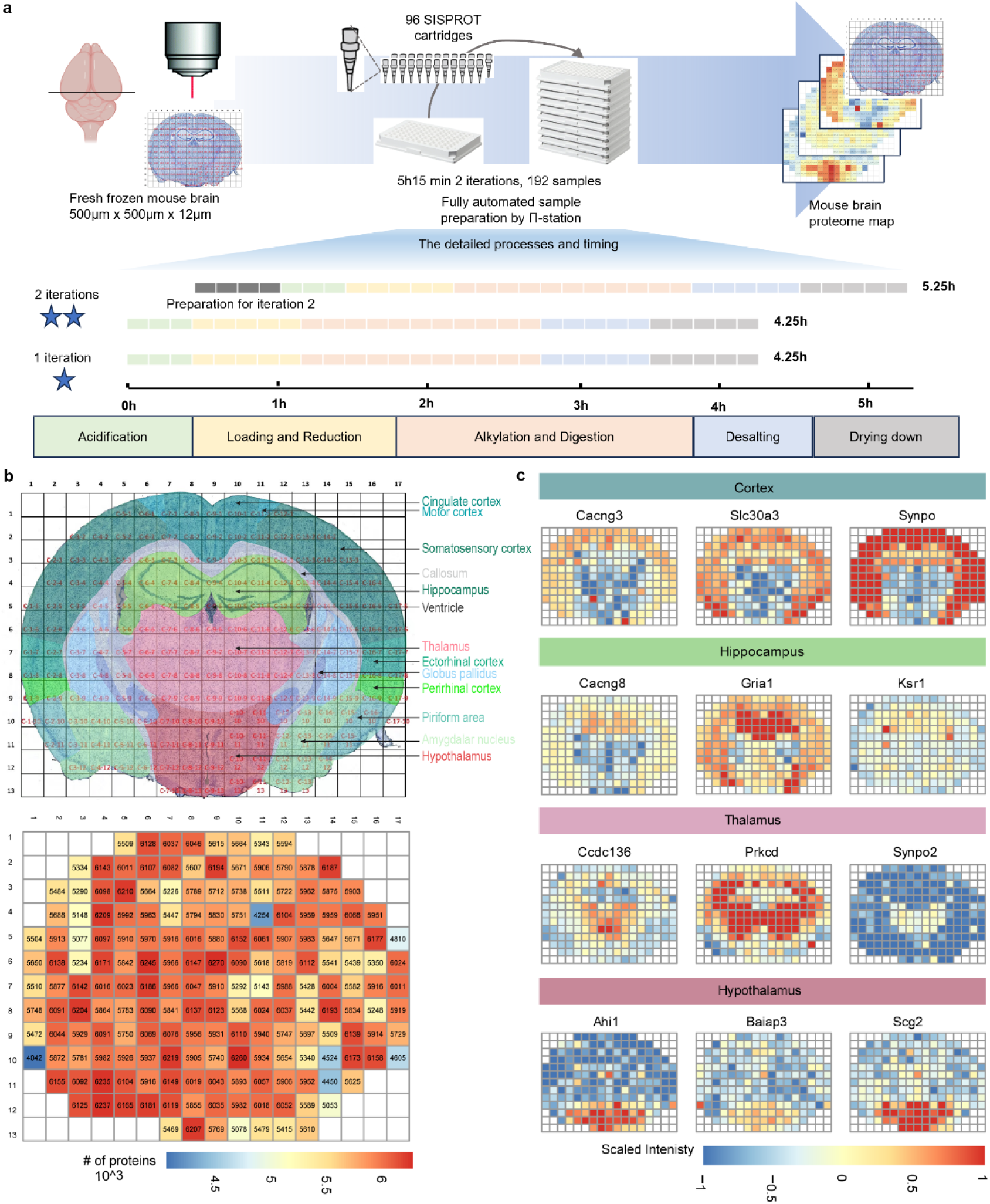
The workflow of spatial proteomic analysis at the π-HuB data factory and the results of the spatial mouse brain proteome. **a,** Schematic of the spatial proteomic workflow. **b,** Schema of the coronal section of the mouse brain and the number of proteins identified in each micro-specimen. The color scale ranges from 4 (blue) to 6.5 (red). **c,** The maps of region marker proteins, which correlated well with the literature. The color scale is from −1.0 (blue) to 1.0 (red).

In summary, we developed the π-Station, a fully automated proteomic analysis platform equipped with feedback control systems. This platform enables ultra-high-throughput generation of high-quality proteomic data from microscale samples with minimal human intervention, greatly increasing productivity and efficiency while reducing errors and costs. It is operating as a core component of the π-HuB data factory, a key pillar of the π-HuB project. Up to date, it has enabled the generation of more than 40,000 high-quality proteomic datasets, supporting dozens of studies funded by the pilot initiatives of π-HuB. Considering that a large monetary investment is required, this platform, at least in its current format, might be cost-prohibitive for individual academic laboratories or core facilities built at most research institutions. Nonetheless, the informatics pipeline for feedback control of systems performance can be easily adopted by any proteomics laboratory right away, allowing prompt maintenance of the LC-MS/MS system and implementing a stop-loss strategy immediately. We have applied it at the proteomic core facility at the National Center for Protein Science in Beijing. Furthermore, the flexible nature of this platform allows for future adaptations to various π-HuB pursuits (e.g., post-translational modifications, protein-protein interactions, and perturbation experiments, and so forth), although it is currently primarily applied to global proteome profiling of biofluids and tissues. Our future endeavors using this platform will mainly focus on producing multi-layer proteomics data with those samples from the π-HuB project. Meanwhile, we are now developing a more compact and cost-effective platform to facilitate the establishment of unmanned proteomics laboratories worldwide for π-HuB and beyond.

## Methods

### Mouse tissue collection

All mice used in this study were bred and housed under institutionally approved conditions in the Animal Experiments Center at Southern University of Science and Technology. All experimental procedures were conducted following approval from the Institutional Animal Care and Use Committee at Southern University of Science and Technology in China. Mouse brains were collected from 6-to 8-week-old C57BL/6 mice.

### Plasma sample collection

We collected a total of 398 plasma samples from patients and healthy plasma from donors. Whole blood was collected into EDTA-treated tubes. Plasma was obtained by centrifugation at 2,000 x g for 15 minutes within 4 hours of collection. Plasma samples were stored at −80°C until processing. All experiments were conducted with the approval of the Research Ethics Committee and according to the Declaration of Helsinki.

### Cell culture

In the methods development, evaluation, and quality control segments, we used ten different types of cell lines. HEK 293T, HeLa, HepG2, Huh7, MCF7, and MDA-MB-231 cells were cultured in Dulbecco’s Modified Eagle Medium (DMEM; Gibco), supplemented with 10% fetal bovine serum (FBS) and 1% penicillin-streptomycin (Gibco). H1975, SNU475, and PLC cells were cultivated in RPMI-1640 medium (Gibco) supplemented with 10% FBS. Additionally, A549 cells were grown in an F-12K nutrient mixture (Gibco), supplemented with 10% FBS and 1% penicillin/streptomycin. All cells were cultured at 37 °C in a 5% CO2 incubator with 95% air humidity. The cells were harvested when they reached approximately 80% confluence. After harvesting, the cells were washed three times with phosphate-buffered saline (PBS) and then stored at −80 °C for future processing.

### Sample preparation of LCM samples

A fresh frozen mouse brain was embedded in Tissue OCT-Freeze Medium (Sakura Finetek USA, Inc.) and sliced into coronal sections with a thickness of 12 μm by using a Leica CM 1900 cryostat (Leica) at −20 °C. The sections were flat mounted onto membrane-coated glass slides (2.0 μm, PEN-membrane, Leica) and fixed by ice-frozen methanol for 10min. Then, the fixed sections were subjected to hematoxylin-eosin staining (Servicebio) and dehydration via an ethanol series. After being scanned using a Leica DM 2500 microscope at 5× magnification, the coronal sections were dissected into spots with a length of 500μm and a width of 500μm using an LMD7 Laser Microdissection Microscope (Leica). Dissected tissue sections were collected into 0.2 mL microtubes (Axygen) were lysed in 50μL of lysis buffer (1% DDM, 10mM HEPES, 150mM NaCl, 600mM guanidine HCl, and 1% Roche protease inhibitor mixture, at pH 7.4) via ultrasonication by a Q800R3 minsonicator (Qsonica) for 15 min (20s on, 20s off, 85% amplitude) at 4 °C. Then the samples were heated in a metal bath at 95 °C for 60 min. After centrifugation, the supernatant was transferred to a new 96-well plate for automated sample preparation.

The sample preparation process using autoSISPROT was effectively carried out on the Π-Station with disposable SISPROT-based cartridges using reagents from the SISPROT kit. This workflow consists of 6 essential steps: (1) Acidification: Begin by acidifying the samples with formic acid utilizing the Biomek i7. Add 5μL of Buffer Acidize to the sample and mix thoroughly by pipetting up and down 20 times. (2) Preparation of SISPROT-based cartridges: Activate the SISPROT-based cartridges with 60μL of Buffer Activate, followed by equilibration using 60μL of Buffer Wash on the Bravo system. (3) Sample Loading and Reduction: Load the samples onto the cartridges, followed by washing with 30μL of Buffer Wash and 30μL of Buffer Activate. Next, reduce samples with 30μL of Buffer Induce in the dark for 15 minutes. Cartridges were transferred into a 3D-printed dark box for dark incubation. (4) On-Tip Digestion: Adjust the pH by adding 30μL of Dissolve A to the cartridges, then introduce 12μL of the Digest Mix. Transfer the cartridges in the 3D printed box and then incubate for 80 min at 37°C for 80 min in Inheco. (5) Peptide Desalting and Elution: Following incubation, return the cartridges to the Bravo. Add 60μL of Buffer Transfer, perform two washes with 60μL of Buffer Wash, and then use 60μL of the Elute buffer to achieve efficient peptide transfer, washing, and elution. (6) Drying Peptides: Transfer the elution plate to the CombiDancer and evaporate the eluted peptide solutions in the 96-well plate at 55°C for 42 minutes.

For a detailed overview of this workflow, please refer to the Supplementary Information. A single iteration of the workflow takes 4.25 hours, while two iterations take 5.5 hours. Additionally, each pair of iterations can be grouped together as a block and can be conducted indefinitely by replenishing reagents and labware.

### Sample preparation of plasma samples

This 96-well format workflow for plasma analysis was developed according to the previously published plasma proteome profiling protocol ^16^. It consists of sample lysis, reduced, alkylated, trypsin digestion, peptides purification (desalting), and preparing peptide solutions for LC-MS/MS analysis. In detail:

1. Upload Samples, reagents, and labware (e.g., containers and tips) to Cytomat hotels under the inventory management of Momentum Software.
2. Transfer samples, reagents, and labware to the liquid-handling station: After initiating the process, transfer the peeled sample and buffer plates and other labware to the Biomek i7 by F7 and Spinnaker robot
3. Denature, lyse, reduce and alkylate samples: transfer 4 μL plasma to a new plate, make 1:25 dilution by adding lysis buffer (1% SDC, 40 mM CAA, 10 mM TCEP, 100 mM Tris-HCl, at pH = 8.8), mix thoroughly by pipetting 3 times (30 μL/time) up and down, transfer the plate from Biomek i7 to ATC and heat the plate at 95°C for 5 min at ATC, and then transfer the plate to Covaris for 5 min sonication with 10 s on and 20 s off.
4. Trypsin digestion: Centrifuge the plate at 2000 rpm for 2 min in Rotanta, transfer Xpeel peeled plate to Biomek i7, transfer 20 μL of the diluted plasma into a new plate, add 80 μL of 100 mM Tris-HCl (pH 8.8), add 2 μg of trypsin, mix thoroughly by pipetting 3 times up and down, and transfer the digestion plate to Inheco for 2h digestion at 37 ℃.
5. Peptide desalting: Centrifuge the plate at 2000 rpm for 1 min in Rotanta, transfer the peeled digestion plate to Bravo, add 15 μL of 10% Formic acid to stop the digestion, seal the plate by ALPS3000, and centrifuge at 6500 rpm for 7min, load the supernatant to pre-equilibrated Agilent C18 cartridges and purify peptides via Bravo peptide desalting program with 1% formic acid water and 80% Acetonitrile in 1% formic acid water.
6. Prepare peptide solutions for LC-MS/MS analysis: concentrate the eluted peptides in CombiDancer at 55 ℃ for 42 min, re-suspend the peptide mixture in 30 μL of 0.1% formic acid water, measure the peptide concentration by Pierce Quantitative Colorimetric Peptide Assay kit in Multiskan SkyHigh, adjust peptides to a concentration of 500 ng/μL, transfer the peptide plate either to Vanquish loader for anlysis of Cytomat 2 Hotel for waiting analysis.

For a comprehensive overview of this workflow, please refer to the Supplementary Information. The duration for a single complete iteration of the workflow is 7.5 hours. Each additional iteration requires an incremental time of 1.5 hours. Moreover, this workflow can be sustained indefinitely through the replenishment of necessary reagents and labware.

### Sample preparation of cell line samples

The cell line sample analysis pipeline was developed based on the in-solution digestion method with sodium deoxycholate (SDC). Similar to the plasma pipeline, peptide desalting was performed on the Bravo platform, while other liquid-handling steps were executed using the Biomek i7. This pipeline can flexibly process samples with 500 ng to 200 µg of proteins by simply adjusting several volumes. In this manuscript, we analyzed proteins ranging from 500 ng to 20 µg to demonstrate sensitivity, using 20 µg of protein as the standard input for all cell line analyses. Cells were incubated in SDC lysis buffer (1% SDC, 100 mM Tris-HCl, pH 8.5) at 95 °C for 10 min and sonicated using a Qsonica Q800R3 sonicator for 10 min (85% power, 10 s on, 10 s off) to shear DNA and RNA. Protein concentrations were measured using a BCA assay. Then, TCEP (final concentration of 10 mM) and CAA (final concentration of 40 mM) were introduced, and the mixture was incubated for 10 minutes at 65 °C for reduction and alkylation. The automated pipeline includes the following steps:

1. Upload Samples, reagents, and labware (e.g., containers and tips) to Cytomat hotels under the inventory management of Momentum Software.
2. Transfer samples, reagents, and labware to the liquid-handling station: After initiating the process, transfer the peeled sample and buffer plates and other labware to Biomek i7 by F7 and Spinnaker robot
3. Preparation for digestion: Adjust the volume of lysates to 50μL by adding 100 mM Tris-HCl (pH 8.8), mix thoroughly by pipetting 3 times up and down. For a protein input of 20 μg, add 30 μL of 100 mM Tris-HCl (pH 8.8) to samples that have a protein concentration of 1 μg/μL.
4. Trypsin digestion: add trypsin to samples at an enzyme-to-protein ratio of 20:1, mix thoroughly by pipetting 3 times up and down, and transfer the digestion plate to Inheco for 2h digestion at 37 ℃.
5. Peptide desalting: Centrifuge the plate at 2000 rpm for 2 min in Rotanta, transfer the peeled digestion plate to Bravo, add 50 μL of 100 mM Tris-HCl (pH 8.8) and 15 μL of 10% Formic acid to stop the digestion, seal the plate by ALPS3000, and centrifuge at 6500 rpm for 7min, load the supernatant to pre-equilibrated Agilent C18 cartridges and purify peptides via Bravo peptide desalting program with 1% formic acid water and 80% Acetonitrile in 1% formic acid water.
6. Prepare peptide solutions for LC-MS/MS analysis: concentrate the eluted peptides in CombiDancer at 55 ℃ for 42 min, re-suspend the peptide mixture in 0.1% formic acid in water by Biomek i7 (for samples with 500ng, 1μg and 2μg proteins, add 8 μL; for samples with over 5μg proteins, add 15 μL), measure the peptide concentration by Pierce Quantitative Colorimetric Peptide Assay kit in Multiskan SkyHigh (optional), adjust peptides to a concentration of 500 ng/μL(optional), transfer the peptide plate either to Vanquish loader for anlysis of Cytomat 2 Hotel for waiting analysis.

For a comprehensive overview of this workflow, please refer to the Supplementary Information. The duration for a complete iteration of the workflow is 7.5 hours, and each additional iteration requires an extra 1.5 hours. Moreover, this workflow can be sustained indefinitely by replenishing the necessary reagents and labware. It is also capable of processing lysates from various types of samples, including tissues.

### LC-MS analysis of LCM samples

Samples were measured with a timsTOF pro mass spectrometer (Bruker Daltonics) coupled to an UltiMate 3000 RSLCnano liquid chromatography system (Thermo Fisher Scientific). Dissolved peptides were loaded to a 20 cm analytical column (100 μm i.d., packed with 1.9 μm C18 particles) and separated at a flow rate of 300 nl/min using a 60-min gradient from 5% to 32% buffer B (acetonitrile with 0.1% formic acid). The mass spectrometer was operated in data-dependent acquisition (DIA) mode with 8 dia-PASEF scans separated into 4 ion mobility windows per scan covering an m/z range from 400 to 1040 by 20 Th windows and an ion mobility range from 0.75 to 1.3 V s cm−2. The capillary voltage was set to 1,750 V. The accumulation and ramp time were specified at 166 ms. The collision energy was decreased from 59 Vs at 1/K_0_ = 1.6 V cm^−2^ to 20 eV at 1/K_0_ = 0.6 Vs cm^−2^.

### LC-MS analysis of plasma samples

Samples were analyzed using an Orbitrap Exploris 480 mass spectrometer (Thermo Fisher Scientific) coupled with a Vanquish Neo UHPLC system (Thermo Fisher Scientific). Vanquish UHPLC Loader and Xcalibur sequence were configured in Momentum Workflow Scheduling Software to enable automated LC-MS analysis. Samples were analyzed by 48SPD DIA methods. Peptides were transferred to a 15 cm Kinetex C18 column (300 μm i.d., packed with 2.6 μm C18 particles, Phenomenex) and separated at a flow rate of 9 μL/min using a 26-min gradient from 6% to 40% buffer B (80% acetonitrile with 0.1% formic acid). The mass spectrometer was operated in data-dependent acquisition (DIA) mode. Full-MS scans were acquired at 60,000 resolution from 350 to 1000 m/z with a normalized automatic gain control (AGC) target of 300% and a maximum injection time of 45 ms. 19 variable windows with 1 Da overlap were used. The resolution was set to 30,000 at 200 m/z. The normalized AGC target was set at 1900%, and the maximum injection time was at 54 ms. Normalized HCD energy was set to 28%.

### LC-MS analysis of cell line samples

The automated LC-MS/MS data acquisition was initiated by the Momentum Workflow Scheduling Software with a pre-defined Xcalibur sequence file. Samples were analyzed using a 24 SPD DIA method on Orbitrap Exploris 480 mass spectrometers (Thermo Fisher Scientific) coupled with a Vanquish Neo UHPLC system (Thermo Fisher Scientific). Peptide separations were carried out on a 15 cm Kinetex C18 column (300 μm i.d., packed with 2.6 μm C18 particles, Phenomenex) with a 52-min gradient from 6% to 40% buffer B (80% acetonitrile with 0.1% formic acid) at a flow rate of 9 μL/min. For the full MS experiment, one scan acquired over m/z 400–1003 at a resolution of 120,000 with a normalized AGC target value at 300% and an auto maximum injection time. For the MS/MS experiment, a total of 90 scans (30 scans/cycle) were acquired at a resolution at 30,000 with a normalized AGC target value at 1900% and with auto maximum injection time. Precursor ions were fragmented by HCD with normalized collision energy at 26%.

### Development and setup of π-ProteomicInfo

We developed an automated system for data storage, processing, quality control, and LC-MS status monitoring. The system consists of a monitor, storage, processing, analysis, and feedback module. We refer to this system as π-ProteomicInfo. For raw data storage, we used a commercial intelligent content management platform called Anyshare (AISHU Technology Corp). This platform monitors the raw data file on the LC-MS computers and automatically transfers the data to storage servers as soon as it is generated. Anyshare servers are also storage servers for our processed data and files. The data processing, quality control, analysis, and feedback tools were developed in-house using Python and integrated into three user interfaces: SearchClient, SearchServer, and SendInfo. The scripts can be accessed through GitHub.

SearchClient must be installed on the LC-MS computers. It monitors the LC-MS data generation and sends raw data to computers and servers with proteomic software. It also tracks the instruments’ status and sends the information to the computer installed SendInfo.exe. SearchServer must be installed on computers that run proteomic software such as MaxQuant^17^, Proteome Discoverer, or Spectronaut. Once raw data is received in a predefined location, SearchServer will trigger protein identification and quantification in these software applications. It will also extract quality control (QC) information, which includes the number of MS1 and MS2 spectra, peptides, and proteins, as well as details such as cycle time, peak capacity, full width at half maximum, retention time, and data points. This information will be summarized in a text file named “stat_summary.txt,” and the text file will be sent to the computer with the installed SendInfo.exe. SendInfo can be installed on a data processing computer or a separate one. It sends messages about QC and LC-MS status to specialists on WeChat or by email to keep them informed.

For processing data from a large cohort, we prefer using the Linux version of DIA-NN. The automation of protein quantification is accomplished through SearchClient and the command line. SearchClient monitors the generation of DIA raw data and sends this data to the server where DIA-NN is installed. Once an individual run is received, DIA-NN begins processing the data using pre-defined parameters and generates a .quant file. The raw files generated by Thermo Scientific mass spectrometers are first converted to .mzML format automatically before being processed by DIA-NN. After processing individual runs, when the number of .quant files reaches the total sample number of the cohort, DIA-NN aggregates the information from all the .quant files, performs cross-run steps, and calculates the final quantities. Using SearchClient and SearchServer can also achieve large numbers of DIA data analyses; it is an optional method. However, as Spectronaut is time-consuming and prone to crashing, we usually don’t use it for large-scale analysis without request.

### Data processing by π-ProteomicInfo

We use Proteome Discoverer (v2.5) for daily DDA QC of Thermo Scientific LC-MS instruments, MaxQuant (v2.0.3) for Bruker LC-MS instruments, and Spectronaut (v15.7) for daily DIA QC. We implemented all these proteomic software in π-ProteomicInfo. To set everything up, we need to specify a few key locations. First, provide the file location of data acquisition (Acquisition dir) and the temporary file location for the file transfer (Data storage) on SearchClient. Next, on SearchServer, select the type and version of software, the file location for the received directory (Received.dir), and the parameter file. The automated data processing is ready and processes new data once it is generated.

All the results shown in this study are generated by DIA-NN (v 1.8.1)^18^, as it is much faster. For DIA-NN, we only need to choose the file location of data acquisition (Acquisition dir) and the temporary file location for the file transfer (Data storage) on SearchClient. Once the data is received by the server with DIA-NN, the process is initiated by the command line. The analyses were then conducted in a library-free mode with some modifications: methionine oxidation and protein N-terminal acetylation were set as the variable modifications, the maximum number of variable modifications was limited to five, the MBR feature was activated, and cross-run normalization was disabled.

### Data analysis

Data analysis was conducted using R software (version 4.0.3) with the output from DIA-NN in the report titled “pg_matrix.tsv.” In counting the number of peptide or protein identifications for each sample, only those peptides or proteins with valid intensities greater than zero were included. For quantitative analysis, the data underwent log10 transformation and were normalized through median centering. Proteins that exhibited more than 50% missing values in each group were filtered out.

The remaining missing values were addressed using the K-nearest neighbors (KNN) imputation method. Principal Component Analysis (PCA) was carried out on proteins quantified across all experiments involving different cell lines, utilizing the “prcomp” function in R. Gene set enrichment analysis was performed using hallmark gene sets from the Molecular Signature Database (MSigDB) ^19^ with the clusterProfiler package (version 4.10.1)^20^. Figures were created with the ggplot2 and pheatmap packages. We also integrated fundamental data analysis in SearchServer. It enables automated extraction of QC information and generation of plots from quantitative files outputted by MaxQuant, Proteome Discover, and Spectronaut. We mainly use this function of AutoSearch to evaluate the QC data and a single run of cohort samples.

## Data availability

The mass spectrometry proteomics data have been deposited in the ProteomeXchange Consortium (http://proteomecentral.proteomexchange.org) via the iProX partner repository^21^ with the dataset identifier PXD064080. Source data are provided with this paper.

## Acknowledgements

We acknowledge Lichao Yu, Yuhai Du, Wei Luan, and Runsheng Zheng, employees at Thermo Fisher Scientific, for valuable discussion and support. We thank Dr. Nengyuan Hu from Southern Medical University for valuable discussion on the data analysis of the mouse brain. We thank Dr. Chunyan Tian and Xiuyuan Zhang from the PDPM group at the National Center for Protein Science (Beijing) for providing us with the cell lines. We thank Kaixuan Li, Xiuwen Yu, Yeye Leng, Meng Yan, and Jiayi Du for their contributions to the LC-MS/MS data acquisition. We thank all the members of the International Academy of Phronesis Medicine.

This work was supported by the China National Key R&D Program (Grant No. 2021YFA13016000 and 2020YFE0202200), NSFC’s Basic Science Center Program (Grant No. 32088101), the Chinese Academy of Medical Sciences Innovation Fund for Medical Sciences (CIFMS) (2019-I2M-5-063), and the Major Project of Guangzhou National Laboratory (grant NO. GZNL2024A03001). And the Big-Science Infrastructure of Phronesis Medicine, of which the pilot phase is funded by the Guangzhou Development District.

## Supplementary information

Illustration of the process for generating spatial proteome, plasma proteome, and cell proteome data.

## Source data

For Figs. 2 and Extended Data Figs. 5, 7, 8, 9, 10.

## Contributions

F.H., D.W., R.T., and L.T. conceptualized the study. D.W. performed most of the experiments and data analysis. S.W., M.P. contributed to set up the π-Station. C.H., X.H., and L.X. contributed to developing π-ProteomicInfo. X.W., Y.L., K.Z. and Y.X. contributed to the workflow and proteomic experiments of spatial samples. W.C. contributed to the laser capture microdissection of the mouse brain. Y.X. and Y.L. contributed to the workflow and proteomic experiments of plasma samples. Y.X. and B.Q. contributed to the workflow and proteomic experiments of the cell line samples. C.H. contributed to the data analysis. D.W., J. Y., and F.H. wrote the manuscript. All authors read, edited, and approved the final version of the manuscript.

---

**Extended Data Fig. 1.**
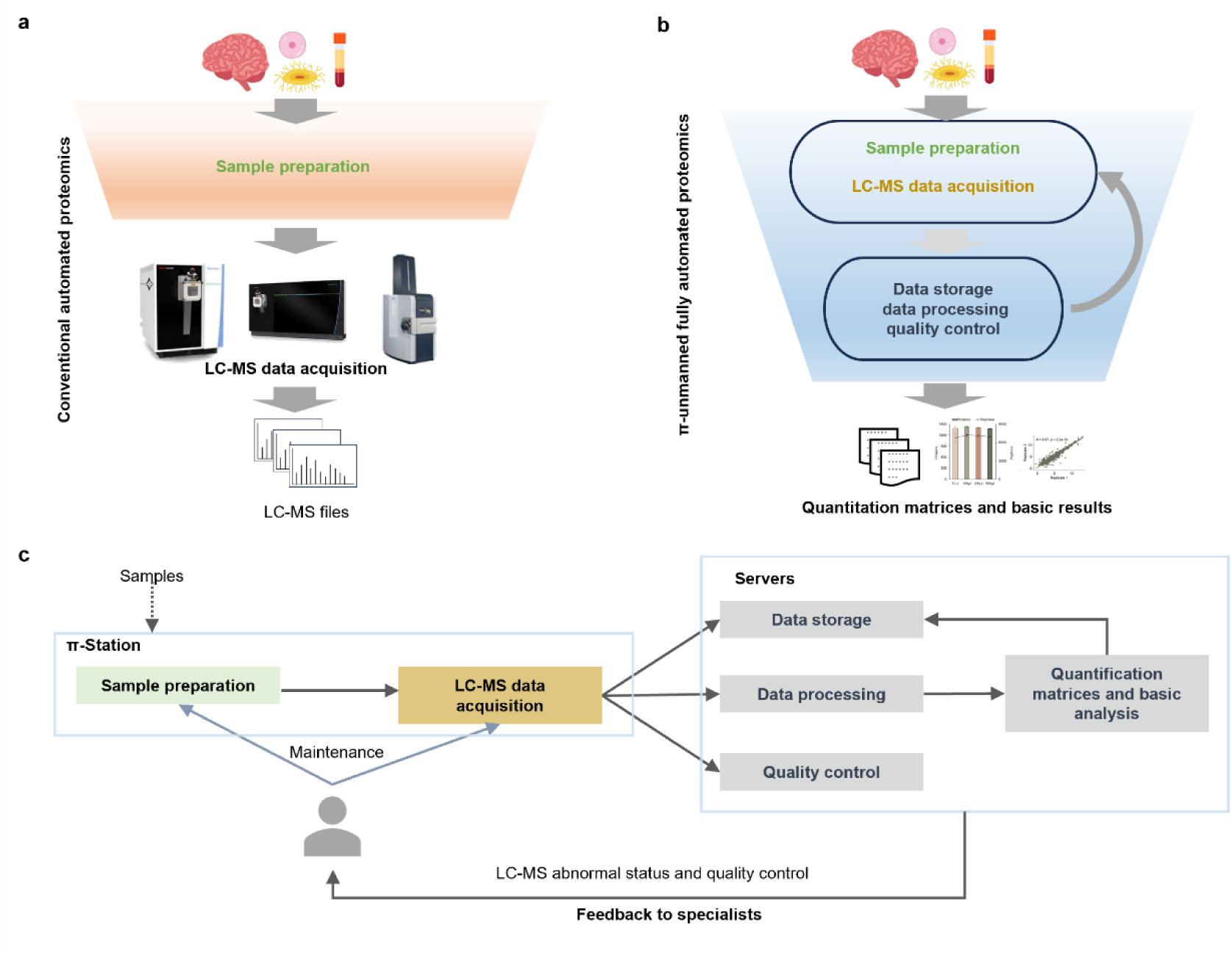
Functional overview of the π-HuB data factory. **a,** Diagram of the conventional automated proteomics, which mainly automates the sample preparation processes and needs human intervention between steps. **b,** Diagram of the π −HuB unmanned fully automated proteomics, which encompasses complete automation of the entire sample-to-data chain with minimal need for human intervention. **c,** The flowchart of the π-HuB data factory. Samples are loaded into the π-Station, where they are automatically prepared into peptides and subjected to LC-MS/MS analysis. The data is automatically stored, processed, and undergoes quality control. If any abnormalities are detected and once quality control information is extracted, the system will notify specialists via a social app or email to facilitate prompt maintenance and prevent sample loss.

**Extended Data Fig. 2.**
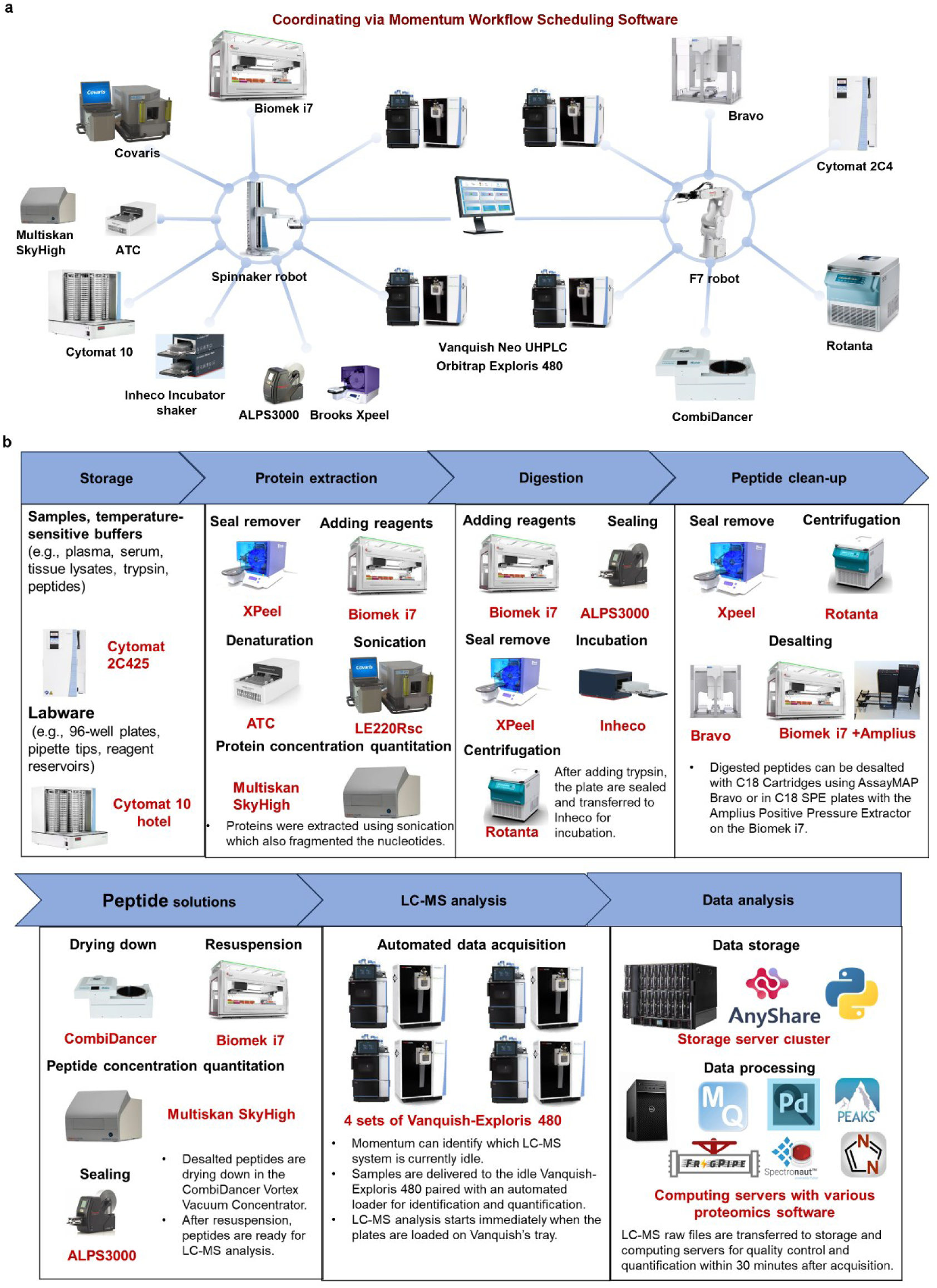
Devices and software of π-HuB data factory. **a,** Customized devices that form the linear, expandable, and connectable π-Station. All these devices are interconnected through the Spinnaker or F7 robot, managed by Momentum software. **b,** Diagram of a typical proteomic analysis workflow, highlighting the devices and software involved at each step of the process.

**Extended Data Fig. 3.**
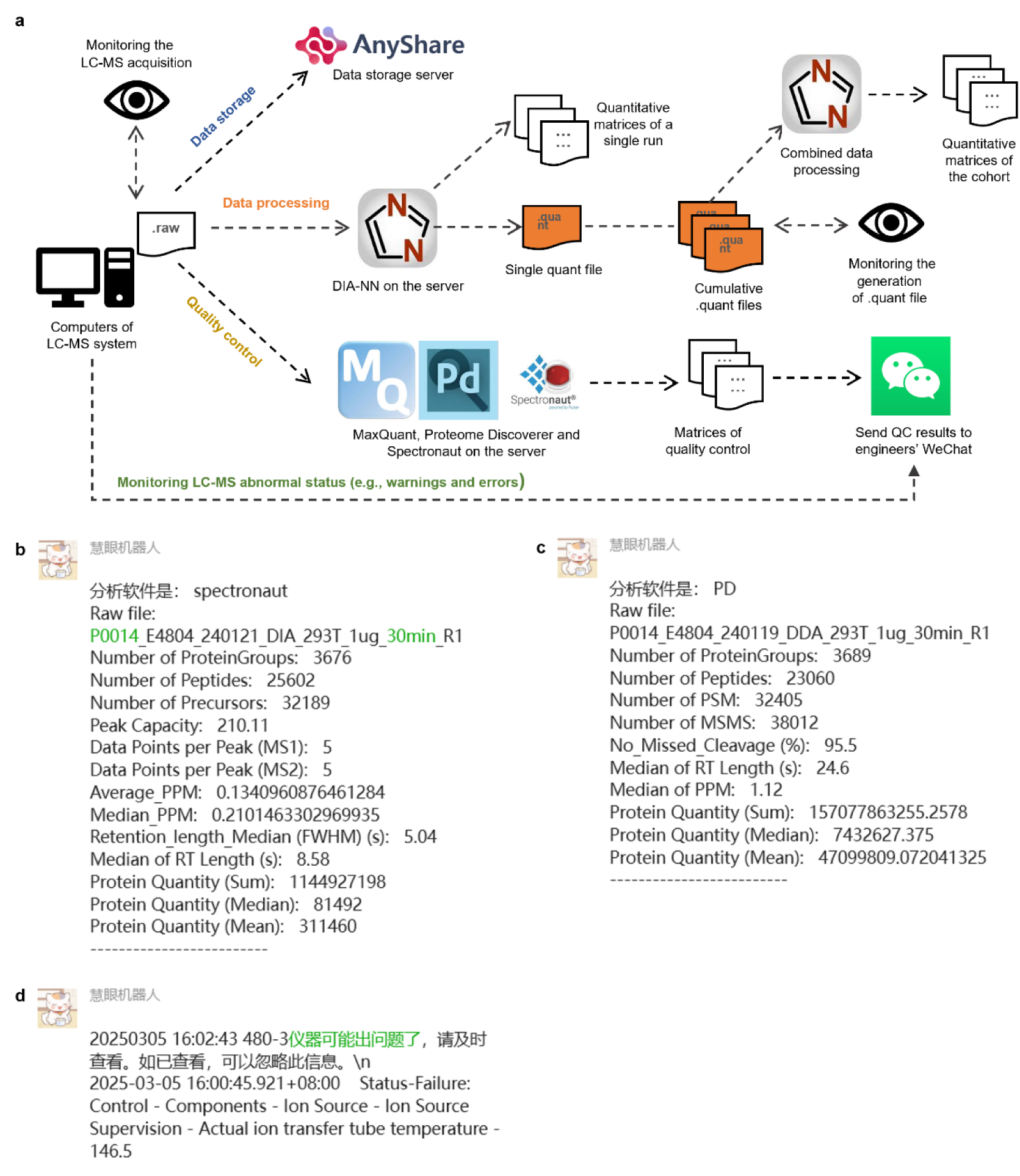
Illustration of the π-ProteomicInfo system. **a,** The workflows involved in data storage, data processing, quality control, monitoring, and feedback within π-ProteomicInfo. **b,** An example of DIA quality control that π-ProteomicInfo extracted and sent to a specialist’s social media application. **c,** An example of DDA quality control that π-ProteomicInfo extracted and sent to the specialist’s social media application. **d,** An example of an abnormal status detected in an LC-MS/MS that π-ProteomicInfo detected and sent to a specialist’s social media application to prompt timely checks and maintenance.

**Extended Data Fig. 4.**
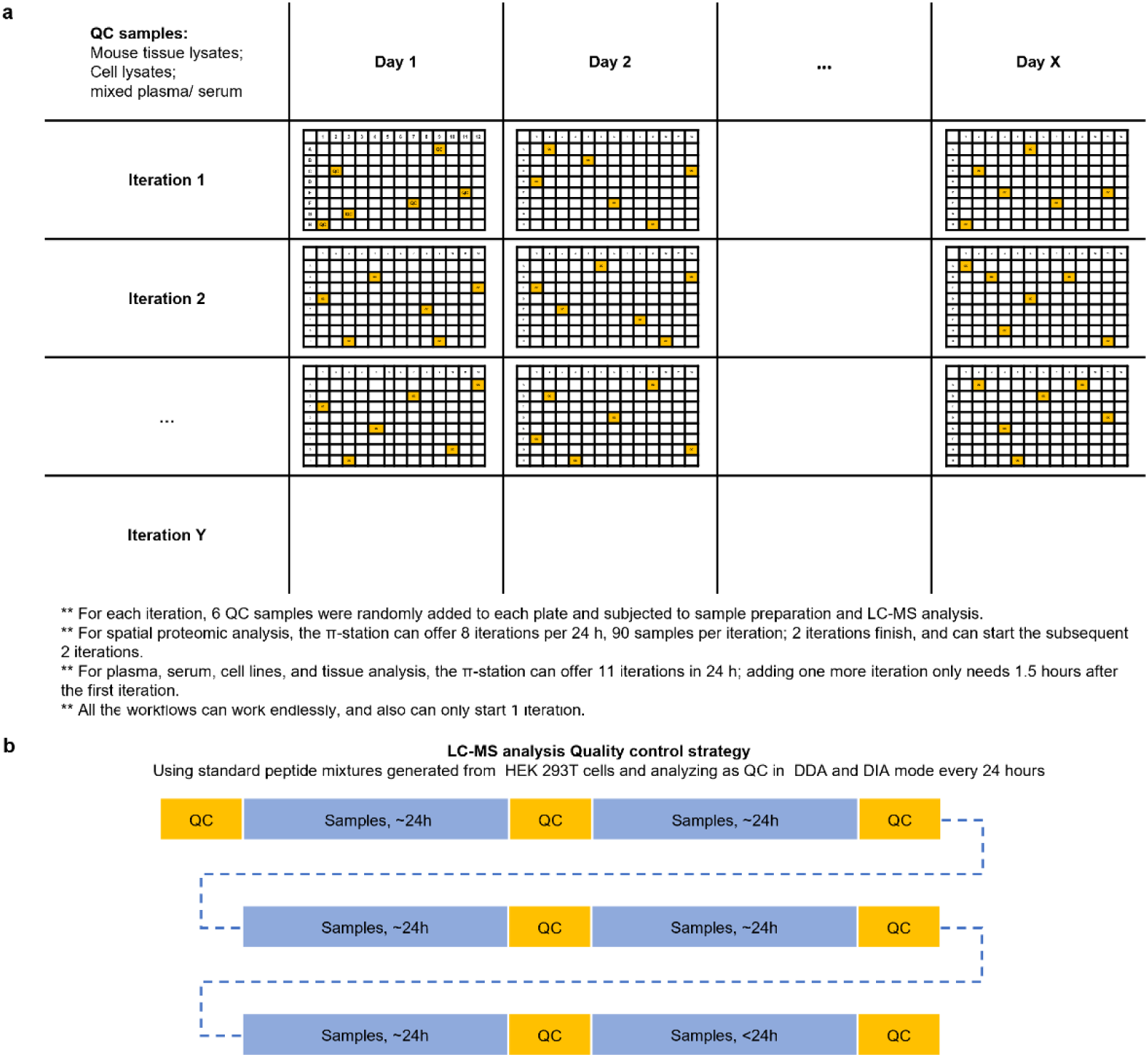
Illustration of the quality control strategies at the π-Hub data factory. **a,** Quality control strategy for sample preparation at π-Station. **b,** Quality control strategy for LC-MS/MS data acquisition.

**Extended Data Fig. 5.**
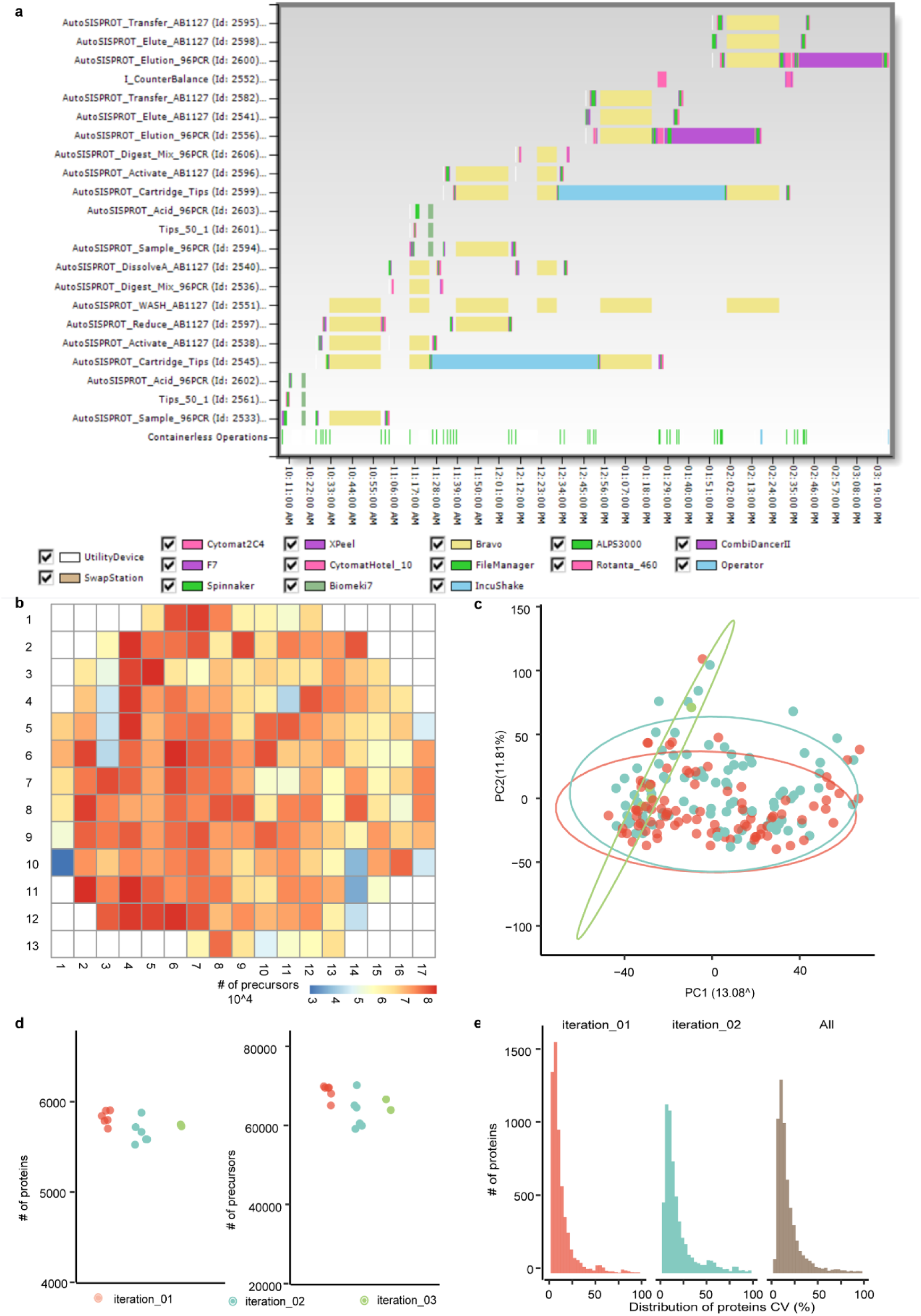
Performance of the spatial proteomic analysis at the π-Hub data factory. **a,** Gantt chart for the spatial proteomic workflow that involves 2 iterations of analysis. **b,** The number of precursors identified in each micro-specimen of the coronal section of the mouse brain. **c,** The principal component analysis (PCA) of samples analyzed across 3 iterations. **d,** The number of proteins and precursors identified in sample preparation quality control. 250 ng of mouse brain lysates were used as the quality control. **e,** The distribution of CVs of protein intensities in sample preparation quality control samples by iterations.

**Extended Data Fig. 6.**
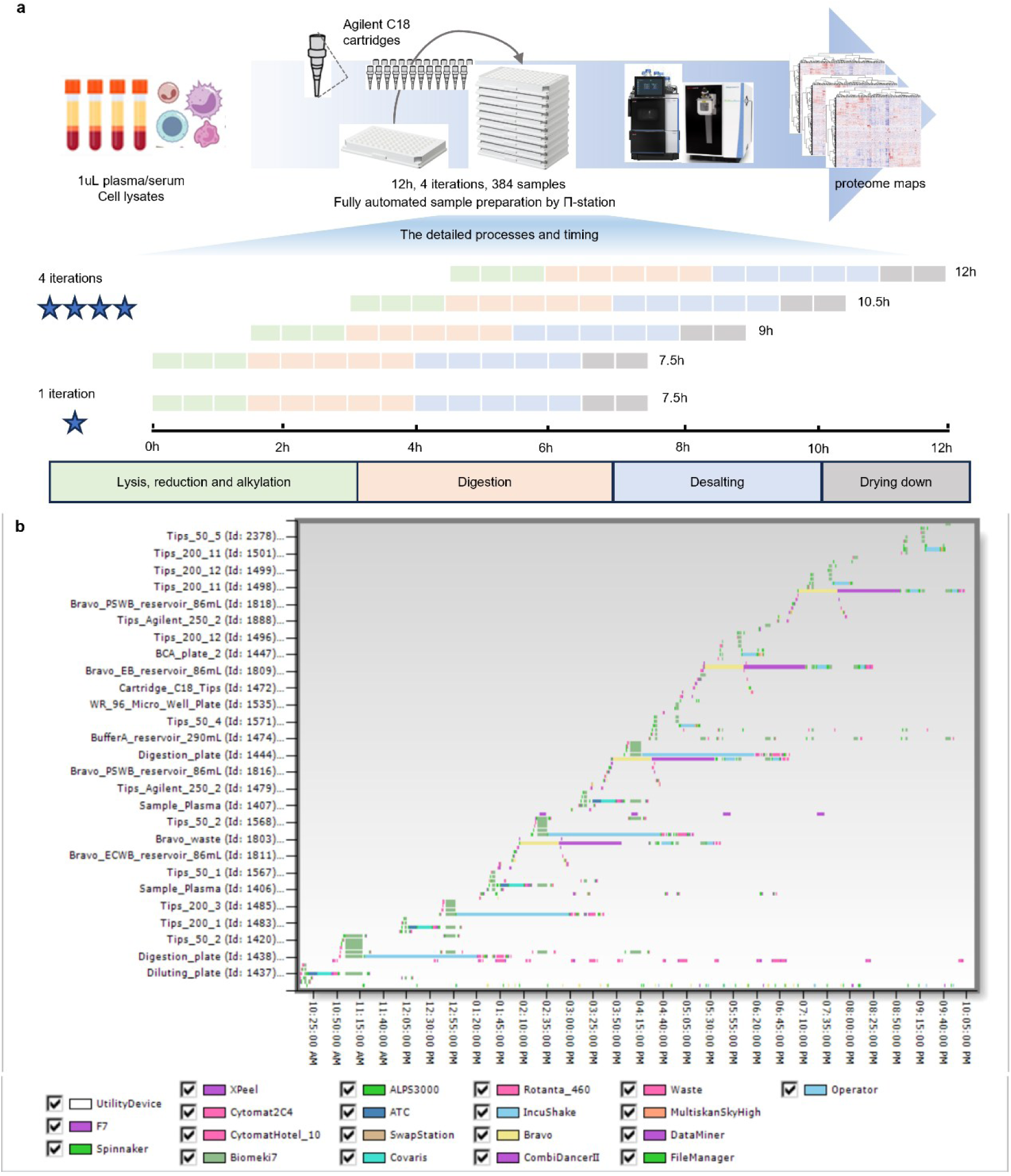
Workflows for the plasma and cell line proteome analysis at the π-Hub data factory. **a,** Schematic of the workflows for analyzing plasma and cell line proteome. **b,** Gantt chart of plasma proteome workflow. Since the workflows for these two sample types are very similar, we have not provided a separate chart for the cell line samples. Detailed descriptions of both workflows can be found in the Methods section.

**Extended Data Fig. 7.**
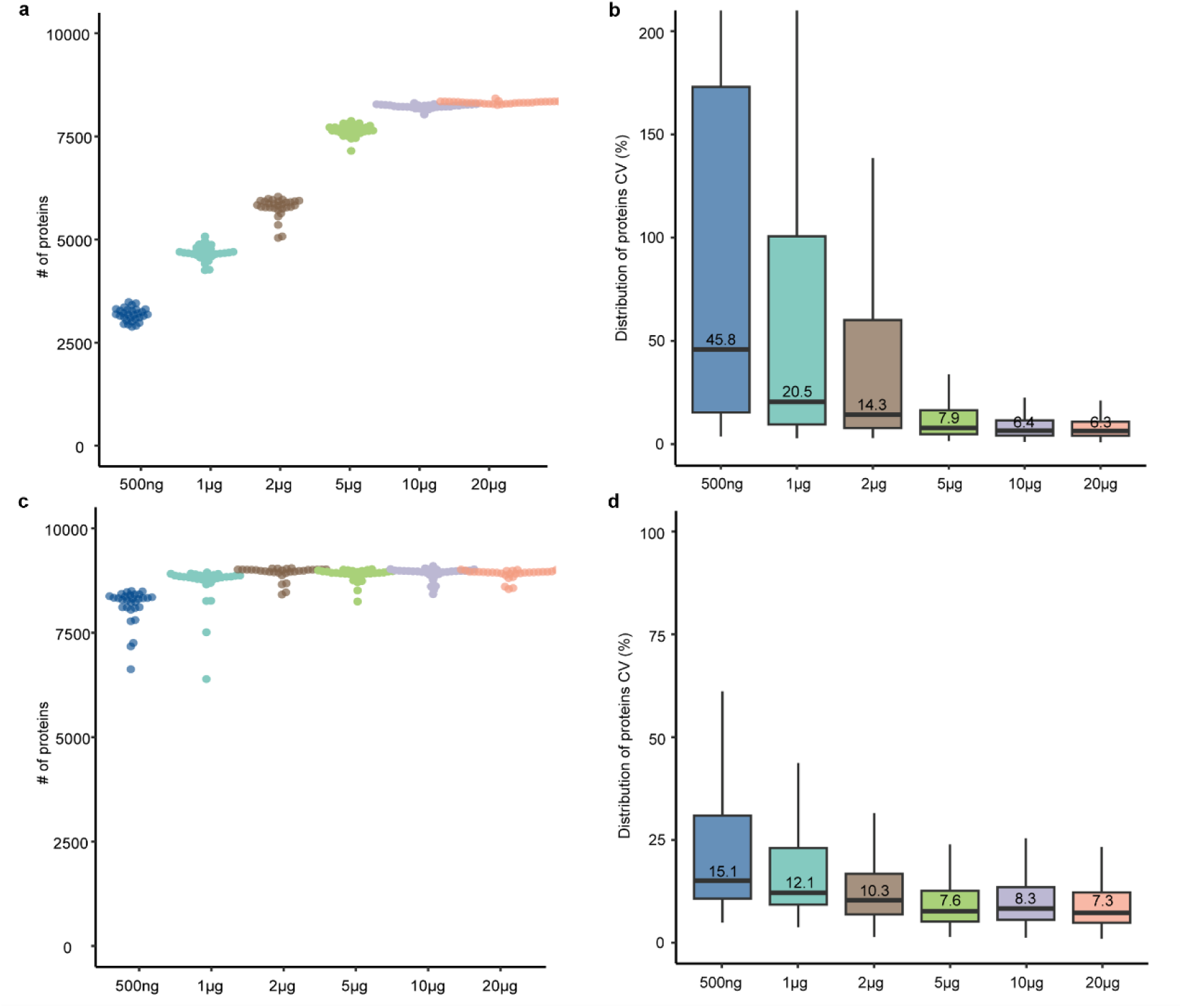
Performance of the π-Hub data factory using different amounts of 293T protein as inputs. **a,** The number of proteins quantified from 500 ng to 20 μg starting amount of protein by the high-throughput LC-MS/MS method. **b,** The distribution of CVs of proteins detected by the high-throughput LC-MS/MS method. **c,** The number of proteins quantified from 500 ng to 20 μg starting amount of protein by the sensitive LC-MS/MS method. **d,** The distribution of CVs of proteins detected by the sensitive LC-MS/MS method.

**Extended Data Fig. 8.**
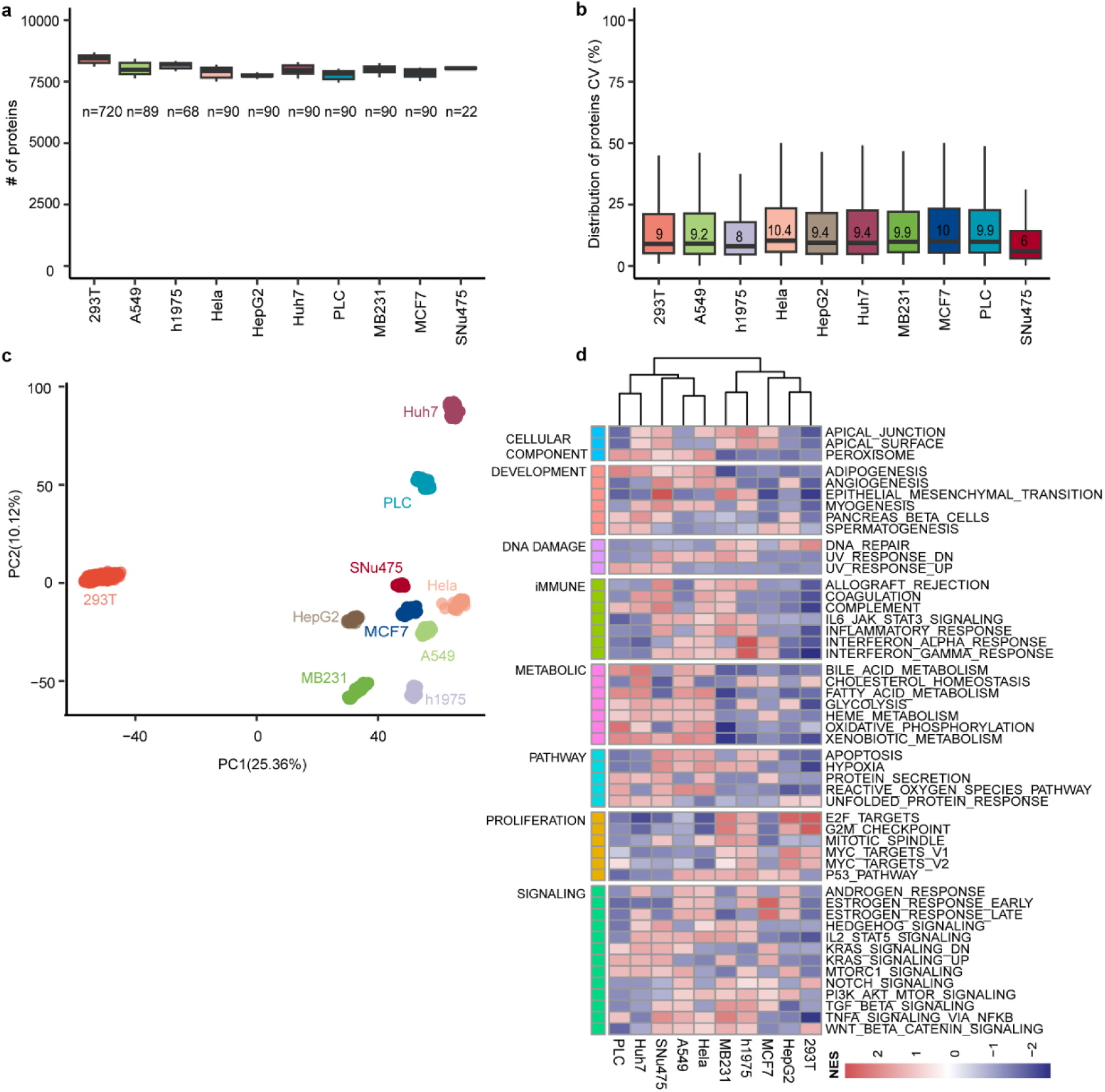
Performance of the π-Hub data factory for analyzing different types of cells. **a,** The number of proteins quantified in each cell line. The number of replicates ranges from 22 to 720. **b,** The distribution of CVs of proteins detected in each cell line. **c,** The principal component analysis of these 10 cell lines. **d,** Single-sample gene set enrichment analysis identifying the biological features of these cell lines.

**Extended Data Fig. 9.**
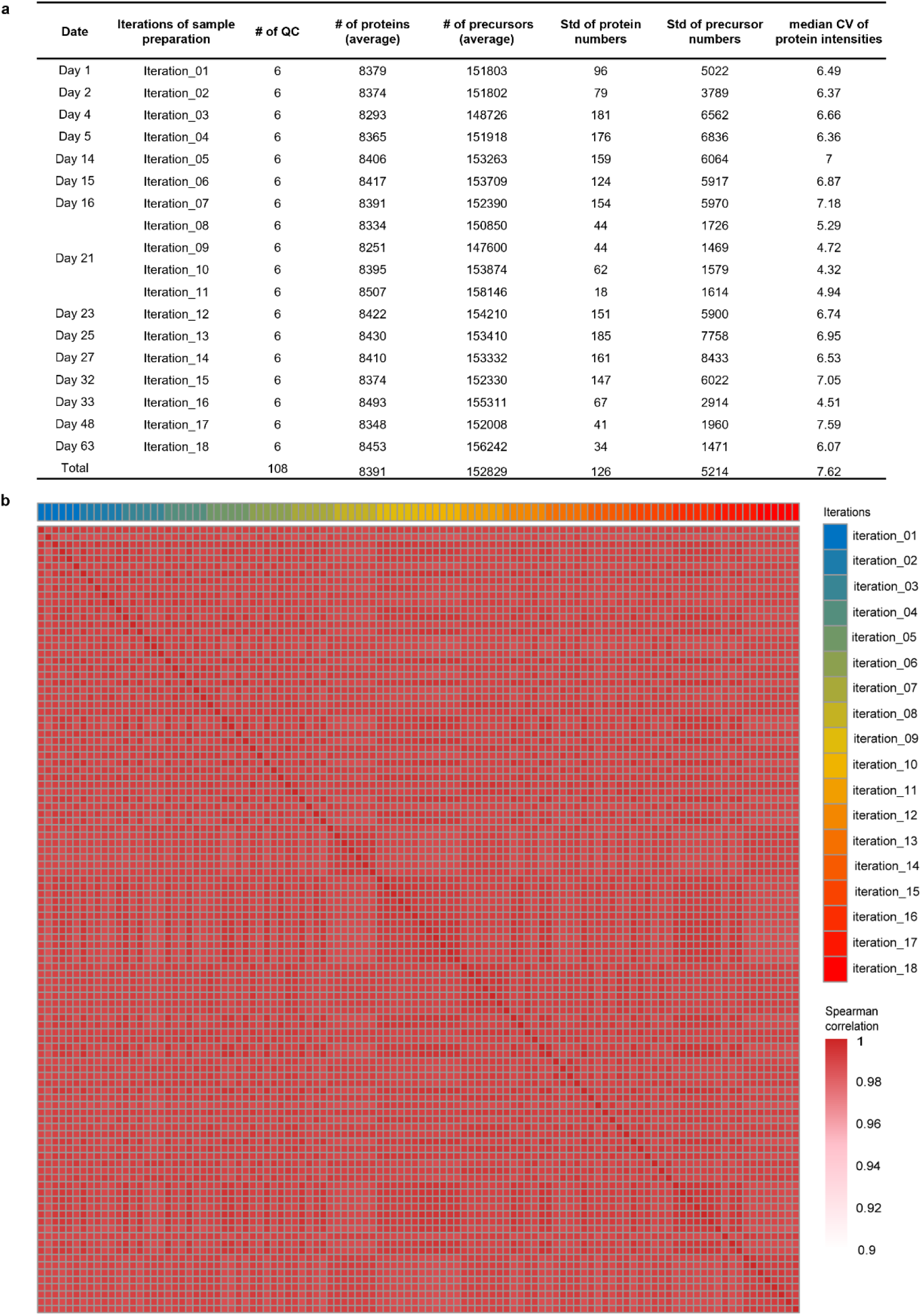
Long-term performance of the π-Hub data factory. **a,** Overview of the quality control data across 18 iterations over a 2-month period. It includes a summary of the number of proteins and precursors, as well as median CV values for individual days and across days. The variability observed is influenced by both sample preparation and LC-MS/MS data acquisition. In each iteration, 6 QC samples were included and processed alongside the other 90 samples. The sample preparation and data acquisition both occurred over the span of 2 months. **b,** The Spearman correlation heatmap of these 108 QC samples.

**Extended Data Fig. 10.**
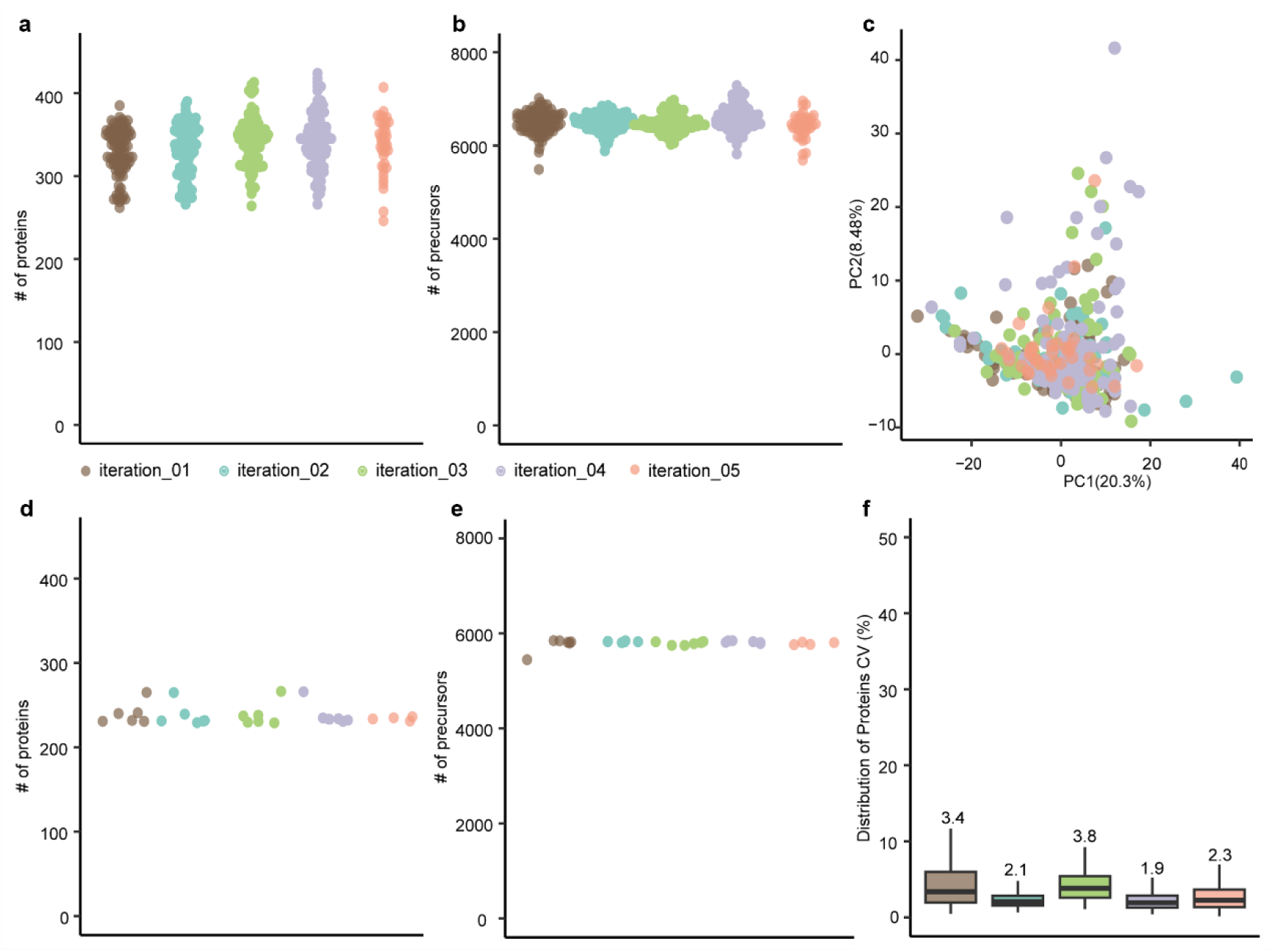
Performance of the π-Hub data factory for analyzing plasma. **a,** The number of proteins quantified in 398 plasma samples, which were processed in 5 iterations. **b,** The number of precursors detected in 398 plasma samples. **c,** The principal component analysis of samples processed in different iterations. **d,** The number of proteins quantified in QC samples. We used a pool of healthy human plasma as QC samples during plasma proteome analysis. **e,** The number of precursors detected in QC samples. **b,** The distribution of CVs for proteins detected in QC samples in each iteration.

## References

1. Bantscheff, M., Lemeer, S., Savitski, M.M. & Kuster, B. Quantitative mass spectrometry in proteomics: critical review update from 2007 to the present. Anal Bioanal Chem 404, 939–965 (2012).

2. He, F. et al. π-HuB: the proteomic navigator of the human body. Nature 636, 322–331 (2024).

3. Hood, L. & Rowen, L. The Human Genome Project: big science transforms biology and medicine. Genome Medicine 5, 79 (2013).

4. Meier, F., Geyer, P.E., Virreira Winter, S., Cox, J. & Mann, M. BoxCar acquisition method enables single-shot proteomics at a depth of 10,000 proteins in 100 minutes. Nat Methods 15, 440–448 (2018).

5. Kawashima, Y. et al. Single-Shot 10K Proteome Approach: Over 10,000 Protein Identifications by Data-Independent Acquisition-Based Single-Shot Proteomics with Ion Mobility Spectrometry. Journal of Proteome Research 21, 1418–1427 (2022).

6. Jiang, Y. et al. Proteomics identifies new therapeutic targets of early-stage hepatocellular carcinoma. Nature 567, 257–261 (2019).

7. Zhang, H. et al. Integrated Proteogenomic Characterization of Human High-Grade Serous Ovarian Cancer. Cell 166, 755–765 (2016).

8. Leutert, M., Rodriguez-Mias, R.A., Fukuda, N.K. & Villen, J. R2-P2 rapid-robotic phosphoproteomics enables multidimensional cell signaling studies. Mol Syst Biol 15, e9021 (2019).

9. Muller, T. et al. Automated sample preparation with SP3 for low-input clinical proteomics. Mol Syst Biol 16, e9111 (2020).

10. Kverneland, A.H. et al. Fully Automated Workflow for Integrated Sample Digestion and Evotip Loading Enabling High-Throughput Clinical Proteomics. Molecular & Cellular Proteomics 23 (2024).

11. Guzman, U.H. et al. Ultra-fast label-free quantification and comprehensive proteome coverage with narrow-window data-independent acquisition. Nature Biotechnology 42, 1855–1866 (2024).

12. Jager, S. et al. In-depth plasma N-glycoproteome profiling using narrow-window data-independent acquisition on the Orbitrap Astral mass spectrometer. Nature Communications 16 (2025).

13. Fu, Q., Murray, C.I., Karpov, O.A. & Van Eyk, J.E. Automated proteomic sample preparation: The key component for high throughput and quantitative mass spectrometry analysis. Mass Spectrom Rev, e21750 (2021).

14. Bubis, J.A. et al. Challenging the Astral mass analyzer to quantify up to 5,300 proteins per single cell at unseen accuracy to uncover cellular heterogeneity. Nature Methods 22, 510–519 (2025).

15. Wu, Q. et al. High-throughput drug target discovery using a fully automated proteomics sample preparation platform. Chemical Science 15, 2833–2847 (2024).

16. Geyer, P.E. et al. Plasma Proteome Profiling to Assess Human Health and Disease. Cell Syst 2, 185–195 (2016).

17. Cox, J. et al. Andromeda: a peptide search engine integrated into the MaxQuant environment. J Proteome Res 10, 1794–1805 (2011).

18. Demichev, V., Messner, C.B., Vernardis, S.I., Lilley, K.S. & Ralser, M. DIA-NN: neural networks and interference correction enable deep proteome coverage in high throughput. Nature Methods 17, 41–44 (2019).

19. Liberzon, A. et al. The Molecular Signatures Database (MSigDB) hallmark gene set collection. Cell Syst 1, 417–425 (2015).

20. Yu, G., Wang, L.G., Han, Y. & He, Q.Y. clusterProfiler: an R package for comparing biological themes among gene clusters. OMICS 16, 284–287 (2012).

21. Ma, J. et al. iProX: an integrated proteome resource. Nucleic Acids Research 47, D1211–D1217 (2019).

